# HDAC inhibitor Givinostat targets DNA-binding of human CGGBP1

**DOI:** 10.1101/2020.01.08.898494

**Authors:** Manthan Patel, Divyesh Patel, Subhamoy Datta, Umashankar Singh

## Abstract

The antineoplastic agent Givinostat inhibits histone deacetylases. We present here our finding that the DNA-binding of human CGGBP1 is also inhibited by Givinostat. CGGBP1, a DNA-binding protein, is required for cancer cell proliferation. In our quest to exploit the potential anti-proliferative effects of CGGBP1 inhibition, we have developed a simple screening assay to identify chemical inhibitors of DNA-protein interactions. We have applied this screen for human CGGBP1 on a library of 1685 compounds and found that Givinostat is a direct inhibitor of CGGBP1-DNA interaction. The mechanism of action of Givinostat should thus extend beyond HDACs to include the inhibition of the myriad functions of CGGBP1 that depend on its binding to the DNA.

## MAIN TEXT

The human CGGBP1 is a repeat-binding 20 KDa protein with widespread expression in normal and cancer cells. It was identified as a transcriptional repressor with cytosine methylation-sensitive DNA binding. More recently, a series of investigations have shown that CGGBP1 is a multifunctional protein (reviewed by Singh and Westermark., 2015 [1]). It regulates transcriptional response to heat shock, undergoes tyrosine phosphorylation upon mitogenic stimulation, and regulates the levels of CDKN1A and TP53 in normal fibroblasts. CGGBP1 localization at midbody prevents tetraploidization. Interestingly, cancer cells (including p53, Rb, p21 or INK4A/ARF loss of function) exhibit a non-oncogene dependence on CGGBP1 similar to the phenomenon described for stress response genes [2]. CGGBP1 depletion increases endogenous DNA damage and activates the G1/S checkpoint.. This involves activation of ATR at telomeres and derepression of the CGGBP1-binding LINEs and SINEs, thereby leading to anomalies in RNA Pol-II activities. CGGBP1 depletion leads to methylation changes at interspersed repeats which could exacerbate the G1/S arrest of cancer cells upon CGGBP1 knockdown. Most interestingly, the normal and cancer cells exhibit different responses to CGGBP1 knockdown: cancer cells undergo G1/S block whereas normal fibroblasts progress through through the S phase slowly arrest in the G2/M phase [3]. Recent works implicate CTCF in regulation of CTCF binding at repeats and regulation of chromatin function [4].

The differential dependence of normal and cancer cells on CGGBP1 suggests that inhibiting it may be a useful means to target cancer cells. A small compound inhibition of CGGBP1 functions may thus synergize the currently employed anti-cancer therapeutic approaches. Since CGGBP1 is a DNA-binding protein, we worked towards identifying a small chemical inhibitor of its DNA-binding. Here we describe a simple and generic library-scale screening method to identify inhibitors of DNA-protein interactions by using CGGBP1 as the target protein of interest. We have pursued some DNA-CGGBP1 interaction inhibitor hits from a widely used library of FDA-approved compounds to identify specific direct inhibitors. Out of 1685 compounds in the library, we have identified Givinostat as a direct inhibitor of CGGBP1 binding to DNA. Givinostat, an anti-cancer agent currently under clinical trials is widely regarded as a specific HDAC inhibitor [5]. Our results expand the scope of mechanisms through which Givinostat acts as an anticancer drug and extend the list of Givinostat targets to include CGGBP1.

## Results and Discussion

Our assay combines the principles of common molecular biology techniques of Southwestern Dot Blot and Immunochemical Detection (abbreviated as DBID). 500 ng of HEK293T genomic DNA (1 kb long fragments) were blotted on nylon membrane discs of 5 mm diameter and crosslinked by vacuum baking at 80°C for 90 minutes (Figure 1A). DNA-linked membrane discs (called dot blots) were blocked and used as probes to capture proteins from cellular lysates. The dot blots were incubated with HEK293T cell lysate to capture DNA-binding protein complexes (Figure 1B). The lysate was pre-incubated with the compounds in the library at a uniform concentration of 100 μM (or 0 μM as a negative control for inhibition) (Figure 1C). The dot blots were incubated with the lysate-inhibitor mix, washed, crosslinked using 4% formaldehyde and probed immunochemically (Figure 1D). For library-scale detection, the dot blots were serially incubated with primary anti-CGGBP1 antibodies, a biotinylated secondary antibody, streptavidin-horseradish peroxidase (HRP) conjugate and a chromogenic HRP substrate. The signals on the dot blots were quantified using densitometry of inverted images of the blots. First, we ensured that the detection was specific by using negative controls. The chromogenic signal detected on the dot blots using the anti-CGGBP1 antibody was specific as the signal was reduced to a weak background when the primary antibody was replaced with isotype-control IgG or serum. Similarly, replacing the lysate with blocking buffer diminished the signal.

**Figure 1.**
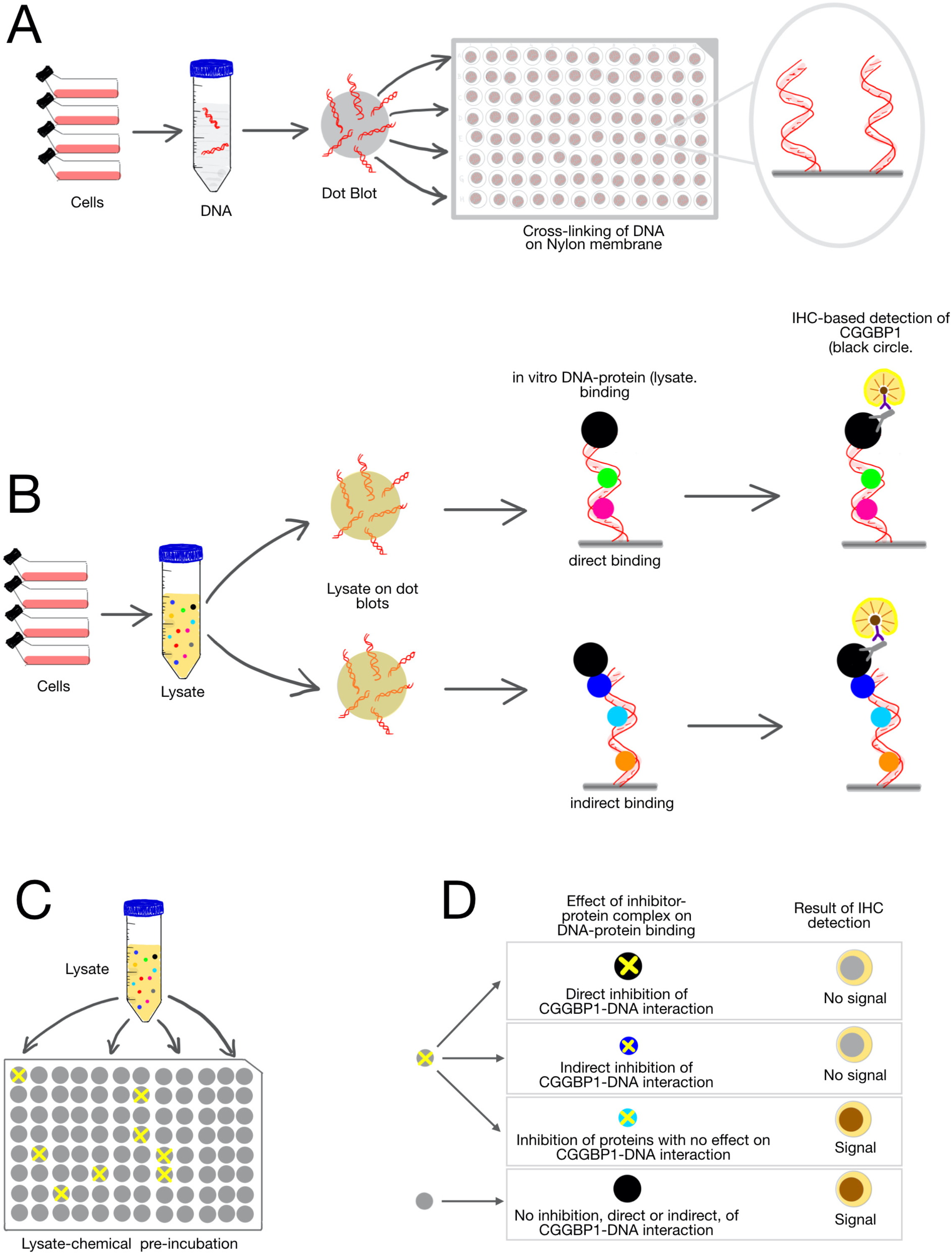
A schematic representation of the **D**ot-**B**lot and **I**mmuno**D**etection (DBID) assay. A: Genomic DNA was isolated from HEK293T cells. Sonicated DNA fragments (mean length of 1kb) were blotted onto positively charged nylon membranes and crosslinked by vacuum heating at 80C. (B) HEK293T cells were lysed and the cleared lysates were used as a source of protein for *in vitro* DNA binding. The assay depicted here is designed to detect binding of CGGBP1 (black circle) to DNA, although this protocol can be generically applied for any protein of interest. CGGBP1 can bind to DNA either directly (top panel) or indirectly (through linker proteins, depicted in blue circle in the bottom panel). The subsequent immunochemistry-based detection reports a brown signal for the direct as well as indirect CGGBP1-DNA complexes alike. The immunochemistry employs a primary antibody against the protein of interest, a biotinylated secondary antibody, streptavidin-HRP conjugates and DAB as the chromogenic substrate and is semi-quantitative in nature. C: Pre-incubation of the cell lysate with inhibitors allows the small molecule compounds to bind to their cognate target proteins in the lysate. Only some of these compounds potentially inhibit the DNA-binding of their target proteins (exemplified with a yellow cross). D: The different possible outcomes of the DBID assay for inhibition of CGGBP1-DNA interactions are depicted. The direct inhibition of CGGBP1 (black circle with a cross) as well as the inhibition of a linker protein (blue circle with a cross) required for CGGBP1-DNA binding are expected to result in “No signal”. Inhibitors that do not have any direct or indirect effect on CGGBP1-DNA interaction show “Signal”.

Pre-incubation with any compound that successfully inhibited DNA-CGGBP1 interactions would result in a weaker signal using anti-CGGBP1 antibody as compared to a mock inhibition (only the compound dilution buffer) (Figure 2A). The lysate was incubated with each compound separately at a final concentration of 100 uM. Majority of the compounds in the library showed no inhibition of CGGBP1-DNA binding, whereas a subset of compounds displayed moderate to strong inhibition of CGGBP1-DNA binding (Figure 2, B and C). Of the 1685 dot blots, one for each compound of the library, the DBID assay returned no inhibition for 1554 compounds, moderate inhibition for 20 compounds, and strong inhibition for 11 compounds (Figure 2). For 100 compounds the assay was inconclusive and we eliminated them from the analysis. The DBID using lysates could not distinguish direct inhibition of CGGBP1 from indirect inhibition.

**Figure 2:**
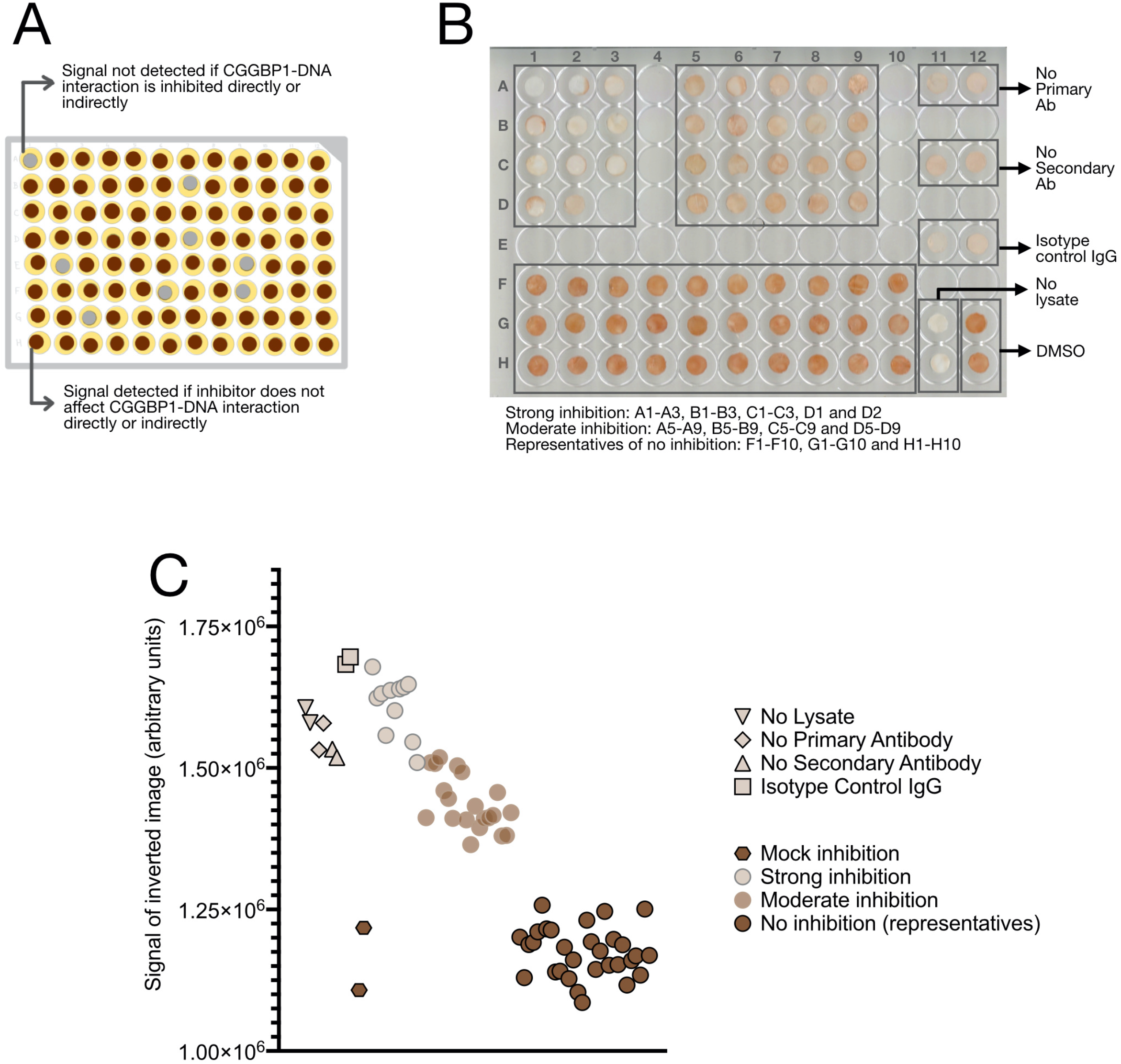
DBID screening of a small molecule chemical library of 1685 compounds identifies inhibitors of CGGBP1-DNA interaction. A: The primary DBID assay was performed in multiple 96-well plates. The lysate was individually pre-incubated with the compounds (one compound per well). After transferring the lysate-compound complexes (as shown in figure 1C) to dot blots (as shown in figure 1A), immunochemical detection was performed using a cocktail of Rabbit anti-CGGBP1 primary antibody. The schematic represents the signals obtained for a 96-well plate. B: The dot blots of the entire library screen for CGGBP1 were manually categorized into 11 strong inhibitors and 20 moderate inhibitors. The actual images of these two groups of dot blots are shown here along with the positive and negative controls as indicated. The identities of the inhibitors are as follows: Strong inhibitors [A1-Givinostat (ITF2357), A2-LRRK2-IN-1, A3-Peficitinib (ASP015K, JNJ-54781532), B1-Ispinesib (SB-715992), B2-TWS119, B3-Domperidone, C1-Gallamine Triethiodide, C2-Moxifloxacin HCl, C3-Sirtinol D1-NAD+, D2-Palbociclib (PD-0332991) HCl], Moderate inhibitors [A5-VX-661, A6-BRD73954, A7-NCT-501, A8-Tenovin-1, A9-Prasugrel, B5-U0126-EtOH, B6-Foretinib (GSK1363089), B7-JNJ-7706621, B8-CHIR-99021 (CT99021), B9-Asenapine maleate, C5-Ethylparaben, C6-LY2874455, C7-Golgicide A, C8-PD173955, C9-Mirin, C5-Ramelteon, C6-Cilnidipine, C7-Dopamine HCl, C8-VR23, C9-AZD3759]. Majority of the compounds in the library did not show any inhibition of CGGBP1-DNA interaction. Thirty representative dot blots of the non-inhibitors are shown in well numbers F1-F10, G1-G10, H1-H10. C: The signals for the dot blots shown in B are quantified by densitometry. The graph shows the signals of the inverted images for each dot blot.

To identify the direct inhibitors of CGGBP1-DNA binding, we used recombinant CGGBP1 (rCGGBP1) containing a C-terminal FLAG tag and a panel of available inhibitors identified in this primary screen. The dot blots were blocked with blocking buffer as described above for the primary screen. Similarly, instead of the lysates, rCGGBP1 was incubated for 30 minutes with the inhibitors identified from the primary screen. The rCGGBP1-inhibitor mix was incubated with the blocked dot blots and probed with anti-FLAG or anti-CGGBP1 antibodies followed by HRP-conjugated secondary antibody incubation and chemiluminescent signal detection (Figure 3, A and B). Signals were quantified and analysed through densitometry (Figure 3C). The inhibitors used for this secondary DBID assay could be again classified as having (i) low or no detectable direct inhibition (Palbociclib, BRD73954, Peficitinb, LRRK2-IN-1 and Ispinesib) or (ii) moderate inhibition (Sirtinol and Tenovin-I). Only one compound, Givinostat, displayed a strong inhibition of rCGGBP1-DNA interaction (Figure 3D).

**Figure 3:**
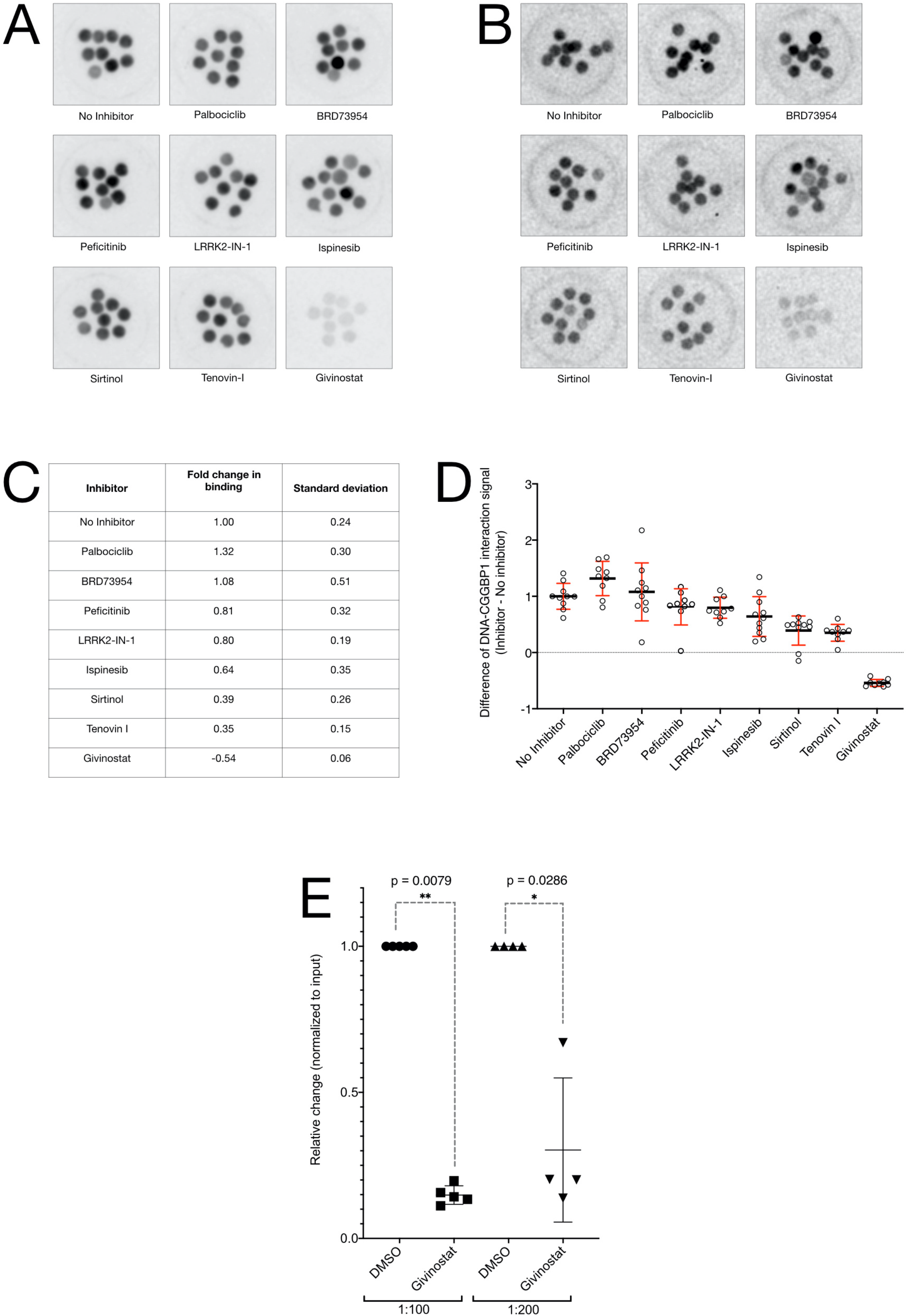
Givinostat acts as a direct inhibitor of CGGBP1-DNA interaction *in vitro*. The secondary DBID screen was performed on a panel of inhibitors identified in the primary screen using FLAG-tagged recombinant CGGBP1 (rCGGBP1). A: rCGGBP1 pre-incubated with compound diluent only (No inhibitor) or the indicated compounds were subjected to DBID by using anti-FLAG antibody. Of the eight different compounds, only Givinostat showed direct inhibition. B: The assay described in A was repeated with anti-CGGBP1 antibody and again Givinostat showed a direct inhibition of DNA-CGGBP1 interaction convincingly. In A and B the multiple dot blots per compound are technical replicates. C and D: Quantification of the signals obtained from the dot blots of secondary screening (B) are tabulated (C) and plotted (D). E: Relative quantification of the amount of Alu DNA immunoprecipitated with rCGGBP1 with or without inhibition with Givinostat was quantified using quantitative PCR and double delta Ct method. The input DNA was used as control and the immunoprecipitated Alu DNA template was used at two different dilutions (1:100 and 1:200). Pre-incubation of rCGGBP1 with Givinostat strongly inhibits the binding of rCGGBP1 directly to pure Alu DNA *in vitro*. Compared to the mock inhibition using DMSO, Givinostat caused a 70-80% reduction in rCGGBP1-Alu DNA interaction.

Finally, we verified the inhibition of rCGGBP1-DNA interaction using an alternative approach. CGGBP1 binds to Alu DNA *in vitro* as well as *in vivo* [1]. We performed *in vitro* immunoprecipitation of Alu DNA using rCGGBP1 with or without inhibition by Givinostat. The difference in the amount of Alu DNA immunoprecipitated using the mock-inhibited rCGGBP1 or Givinostat-inhibited CGGBP1 was quantified using real-time PCR (Figure 3E). Using the amount of input Alu DNA as a control, we could establish that Givinostat is a strong direct inhibitor of CGGBP1-Alu DNA interaction.

The sensitivity of cancer cells to arrest in G1/S transition upon CGGBP1 depletion can be used to retard cancer cell proliferation by using a small molecule inhibitor of CGGBP1. The search for such an inhibitor presents multiple challenges. Conventionally, the small molecule inhibitors of enzyme activities are easier to test because the enzyme assays can provide a direct measurement of the inhibition. Assuming that CGGBP1 is a DNA-binding protein with no enzymatic activity, a similar measurement of CGGBP1 “activity” and the “inhibition of its activity by a compound” is not possible. Thus, we had to first define the “activity” of CGGBP1 and then invent a methodology to screen for inhibitors. Building upon the knowledge that CGGBP1 is a DNA-binding protein, we worked under the constraints of an assumption that DNA-binding of CGGBP1 is required for its functions. As a corollary, an inhibition of DNA-binding of CGGBP1 could be stated as inhibition of CGGBP1. The structure of CGGBP1 is not known and homology modelling does not give reliable structure information for a rational inhibitor design. In the light of the above mentioned challenges, we established the DBID protocol to screen a library of inhibitors. The DBID protocol is simpler than previously described methods [6–8] and can be used for a library level inhibitor screen against any DNA-binding protein for which reliable antibodies are available. Also, by using pure recombinant protein in DBID, chemical libraries can be screened for direct inhibitors.

We identified several indirect and one robust direct inhibitor of CGGBP1. This serendipitous identification of Givinostat is interesting from various viewpoints. It suggests that the antineoplastic activities of Givinostat likely do not depend solely on HDAC inhibition. Although HDACs are a primary target for Givinostat, it also inhibits a mutant form of JAK2 [9,10]. Our results suggest that a certain fraction of the pharmacological effects of Givinostat depend on CGGBP1-inhibition as well. HDAC and CGGBP1 could be inhibited by Givinostat through completely independent mechanisms and it is just that our investigation reveals a potent and useful non-specificity of Givinostat.

## LIST OF ABBREVIATIONS

DBID: 
rCGGBP1: 

## DECLARATIONS

### Ethics approval and consent to participate

Not applicable

### Consent for publication

Not applicable

### Availability of data and materials

An additional file contains details of the methods and materials used.

## AUTHORS’ CONTRIBUTIONS

MP, DP and SD performed the experiments, analyzed the data and participated in writing the manuscript. US provided the supervision, analyzed the data and wrote the manuscript. MP and DP contributed to the work equally. All authors read and approved the final manuscript.

## ACKNOWLEDGEMENTS

The authors acknowledge Dr. Dhiraj Bhatia, Ms. Anjali Rajwar, Dr. Sharmistha Majumdar, Ms. Vasudha Sharma and Dr. Chandrakumar Appayee for collaborations in various experiments important for this work.

## ADDITIONAL FILE

HDAC inhibitor Givinostat targets DNA-binding of human CGGBP1 Manthan Patel, Divyesh Patel, Subhamoy Datta, Umashankar Singh

HoMeCell Lab, Biological Engineering, Indian Institute of Technology Gandhinagar, Gandhinagar, Gujarat. 382355, India.

## Material and Methods

### Cell Culture and Cell lysate preparation

HEK293T cells were grown in DMEM (AL007A, Himedia) supplemented with 10% FBS (RM1112, Himedia) and antibiotic-antimycotic agent (15240062, Gibco). Cells were collected and washed with PBS (TL1006, Himedia). Washed cell pellet was lysed in RIPA buffer (150mM NaCL, 5mM EDTA, 50mM Tris, 1% NP40 (IGEPAL), 0.1 Na-deoxycholate and 0.1 % SDS) containing Halt protease and phosphatase protein inhibitor cocktails (PI78441, Invitrogen).

### Crosslinking of DNA with membrane disc

Genomic DNA was isolated from HEK293T cells. DNA was sonicated using a Diagenode Bioruptor sonicator for 21 cycles at 30 second on followed by 30 second off. Sonication was standardised to get ~ 1kb fragment size DNA. Sonicated genomic DNA was dissolved in 1x TE buffer at 200 ng/µl. Nylon-membrane (GX222020NN, Genetix) discs (5 mm diameter) were soaked with the sonicated DNA overnight. DNA was crosslinked by vacuum heating the wet membranes at 80° C for 90 minutes. DNA crosslinked nylon-membrane discs (called dot blots) were transferred to 96 well plates and incubated with blocking solution (PBST with 10% FBS). For the primary screen one dot blot was used for each of the 1685 compounds (LL100, Selleckchem) distributed across multiple 96 well plates. In addition, the control samples were run on two dot blots for each 96 well plate.

### Dot-blot assay

DNA crosslinked nylon membrane were pre-incubated with blocking solution for 90 minutes. Cell lysate (15 μl of a 25 ml lysate stock obtained from approximately 500 million cells) was incubated with compounds (final concentration 100 uM) or compound diluent (0.05% DMSO in 1xPBS) for 30 minutes. Pre-incubated compound-lysate mix was transferred to DNA cross linked nylon membrane and incubated for 90 minutes at 4° C. Cross linked dot blots were washed three times with PBS and protein-DNA complexes were crosslinked with 4% formaldehyde in 1x PBS. The dot blots were then incubated with 50 μl of rabbit polyclonal anti-CGGBP1 antibody mix (a mix of 1:120 dilutions of SC-292517, SCBT and 10716-1-AP, Proteintech) overnight with gentle rocking. Dot blots were washed with PBST three times and incubated with 30 μl of biotinylated anti-rabbit secondary antibody for 90 minutes (1:10 dilution, 865002, R&D Systems). After three PBST washes the dot blots were incubated with high sensitivity streptavidin conjugated to horseradish peroxidase (1:10 dilution, 865006, R&D Systems) for 90 minutes. All antibody dilutions and streptavidin-HRP conjugate dilutions were done in blocking solution. Dot blots were washed three times with PBST and incubated with 15 μl of DAB (3,3’-Diaminobenzidine) chromogen (2 ml of DAB chromogen (860001, R&D Systems) diluted in DAB chromogen buffer (860005, R&D Systems). Positive and negative controls are described below.

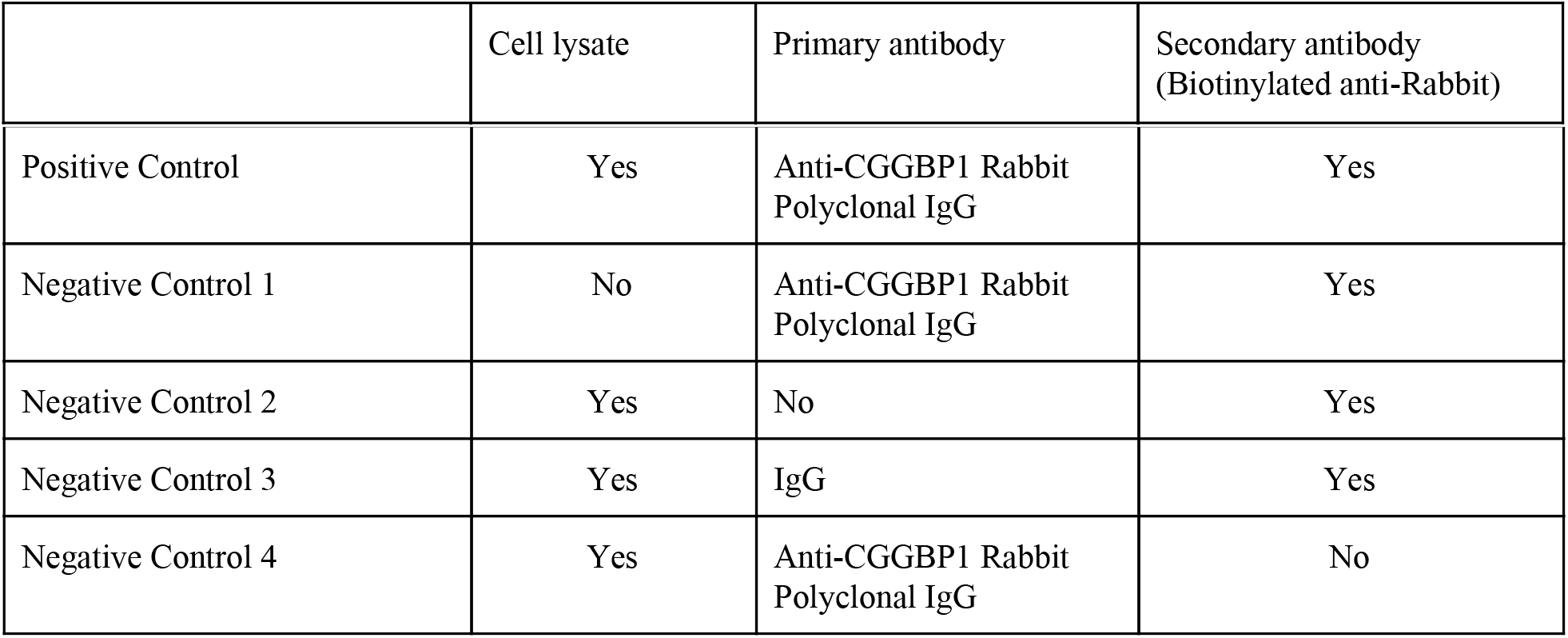

### Secondary screening for direct or indirect inhibition of the primary hits

The secondary screening was performed for eight inhibitors obtained as hits from the primary screen along with a positive control (no inhibitor) and a negative control (no primary antibody). The assay was done in 6-well plates with each inhibitor-rCGGBP1 combination assayed in 8-10 technical replicates. For each sample, dot blots were incubated with 250 μl of blocking solution (10% FBS in PBS) for 1 hour at room temperature in a moist chamber. Simultaneously, the rCGGBP1 (500 ng per compound) diluted in RIPA lysis buffer (containing protease phosphatase inhibitor cocktail) was incubated with inhibitors (at a final concentration of 100 uM for 45 minutes) at 4° C. As a positive control, the compound diluent (0.05% DMSO in 1x PBS). The protein-inhibitor mix was diluted in blocking solution such that the final volume was 250 μl for each sample. The blocking solution was removed and rCGGBP1-inhibitor mix was transferred onto the membranes in each well for 1 hour at 4° C in a moist chamber with gentle rocking. This was followed by fixation of the interactions using 1% PFA for 5 minutes and subsequent PBS washes three times. Membranes were incubated with Mouse anti-FLAG antibody (1:1000 of SC-166384, SCBT, in blocking solution) for 1 hour at room temperature with gentle rocking, followed by washing three times with PBST. The membranes were then incubated with anti-Mouse HRP-conjugated secondary antibody (1:5000 of NA931, GE Healthcare) for 1 hour at room temperature followed by three washes with PBST. The signal was detected with ECL substrate (32106, Pierce). Further the membranes were stripped with 0.2 N NaOH for 5 minutes and incubated with anti-CGGBP1 antibody (1:1000 in blocking solution) overnight at 4° C followed by three PBST washes. The membranes were incubated with anti-Rabbit HRP-conjugated secondary antibody (1:5000 of NA934, GE Healthcare) for 1 hour at room temperature and subsequently washed with PBST thrice. The signal was captured detecting chemiluminescence as described above. The signal was quantified by densitometry analysis of images using ImageJ software. The statistical analysis and data presentation was performed using Open Office and GraphPad Prism8.

### Synthesis of Alu DNA

For the generation of full-length Alu DNA, an established DNA-binding target of CGGBP1, the full length consensus sequence of Alu SINE was synthesized in five overlapping oligonucleotides. The sequences of the oligonucleotides are as follows: The 5’ end of the first fragment and the 3’ end of the last fragment contained the T7 and SP6 primers respectively. The Alu DNA product was obtained through overlapping PCR using an equimolar mix of overlapping oligonucleotides as template and T7 and SP6 sequences as primers. The PCR product was run on the agarose gel and purified using PCR purification kit (A1222, Promega). The Alu PCR product was cloned into using pGEM-T Easy Vector (A1380, Promega). The clones obtained were subjected to Sanger sequencing for verification. This clone was used as a template for amplifying Alu DNA for *in vitro* DNA-rCGGBP1 immunoprecipitation assay. Two different lengths of Alu DNA were amplified due to two priming sites for T7 as well as SP6 in the clone (~320 bp (T7 and SP6 sites in the insert) and other at ~400 bp (T7 and SP6 sites in the vector backbone). The Alu DNA was amplified using either unmethylated cytosine or 5’-methylated dCTPs. The sequence of the Alu DNA is as follows: 5’-TAATACGACTCACTATAGGGGGCCGGGCGCGGTGGCTCACGCCTGTAATCCCAGCACTTTGGGAGGCCGAGGCGGGAGGATCGCTTGAGCCCAGGAGTTCGAGACCAGCCTGGGCAACATAGCGAGACCCCGTCTCTACAAAAAATACAAAAATTAGCCGGGCGTGGTGGCGCGCGCCTGTAGTCCCAGCTACTCGGGAGGCTGAGGCAGGAGGATCGCTTGAGCCCAGGAGTTCGAGGCTGCAGTGAGCTATGATCGCGCCACTGCACTCCAGCCTGGGCGACAGAGCGAGACCCTGTCTCTTCTATAGTGTCACCTAAAT-3’

### *In vitro* DNA-IP and qPCR

The *in vitro* DNA immunoprecipitation was performed with or without Givinostat. rCGGBP1 (0.5 ug) diluted in 1x PBS was incubated with inhibitor (100 uM final concentration) for 45 minutes at room temperature. The final volume was adjusted to 50 μl with PBS. The protein-inhibitor mix was incubated with 1μg of Alu DNA. Simultaneously 3 μg of the anti-FLAG antibody (described above) was subjected to incubation with protein-G sepharose beads (60 ul, 17061801, GE Healthcare) for 1 hour with tumbling. The DNA-protein-inhibitor mix was transferred to the tube containing the anti-FLAG antibody-bound protein-G sepharose and incubated for 60 minutes at room temperature with tumbling. The beads were allowed to settle down followed by gentle spin and the supernatant containing the unbound antibody and DNA was removed. The beads were gently washed three times with ice cold 1x PBS. For each sample the 1x TE buffer (40 μl per sample) was added and mixed followed by heating at 80° for 20 minutes to elute the bound DNA.

### qPCR

The immunoprecipitated Alu DNA in presence and absence of Givinostat was used as template for the qPCR (1725124, Biorad) using T7 and SP6 primers. The PCR was performed for both the samples (Alu DNA immunoprecipitated with Givinostat-inhibited or and mock-inhibited rCGGBP1). The template was used at different dilutions of the immunoprecipitated DNA (1:100 and 1:200 diluted) for qPCR in multiple replicates. The input Alu DNA template was used as a control to calculate the first delta Ct (dCt). The second delta Ct values were calculated by subtracting the dCt values obtained for the mock-inhibited sample from those of the Givinostat-inhibited sample. Following are the PCR conditions for Alu PCR: 95° C-5 minutes, (95° C-20 seconds, 55° C-20 seconds, 72° C-30 seconds, 80° C-30 seconds) x50, melting stage.

## REFERENCES

1. Singh U, Westermark B. CGGBP1--an indispensable protein with ubiquitous cytoprotective functions. Ups J Med Sci. 2015;120:219–32.

2. Nagel R, Semenova EA, Berns A. Drugging the addict: non-oncogene addiction as a target for cancer therapy. EMBO Rep. European Molecular Biology Organization; 2016;17:1516.

3. Singh U, Roswall P, Uhrbom L, Westermark B. CGGBP1 regulates cell cycle in cancer cells. BMC Mol Biol. 2011;12:28.

4. Patel D, Patel M, Datta S, Singh U. CGGBP1 regulates CTCF occupancy at repeats. Epigenetics Chromatin. 2019;12:57.

5. Tan J, Cang S, Ma Y, Petrillo RL, Liu D. Novel histone deacetylase inhibitors in clinical trials as anti-cancer agents. J Hematol Oncol. 2010;3:5.

6. Alonso N, Guillen R, Chambers JW, Leng F. A rapid and sensitive high-throughput screening method to identify compounds targeting protein–nucleic acids interactions [Internet]. Nucleic Acids Research. 2015. p. e52–e52. Available from: http://dx.doi.org/10.1093/nar/gkv069

7. Voter AF, Manthei KA, Keck JL. A High-Throughput Screening Strategy to Identify Protein-Protein Interaction Inhibitors That Block the Fanconi Anemia DNA Repair Pathway. J Biomol Screen. 2016;21:626–33.

8. Chan LL, Pineda M, Heeres JT, Hergenrother PJ, Cunningham BT. A general method for discovering inhibitors of protein-DNA interactions using photonic crystal biosensors. ACS Chem Biol. 2008;3:437–48.

9. Guerini V, Barbui V, Spinelli O, Salvi A, Dellacasa C, Carobbio A, et al. The histone deacetylase inhibitor ITF2357 selectively targets cells bearing mutated JAK2(V617F). Leukemia. 2008;22:740–7.

10. Finazzi G, Vannucchi AM, Martinelli V, Ruggeri M, Nobile F, Specchia G, et al. A phase II study of Givinostat in combination with hydroxycarbamide in patients with polycythaemia vera unresponsive to hydroxycarbamide monotherapy. Br J Haematol. 2013;161:688–94.

